# Nanomolar interactions of alpha-synuclein fibrils to tau determined by FCS

**DOI:** 10.1101/2023.04.13.536728

**Authors:** Jennifer Ramirez, Emily Ann Elizabeth Brackhahn, E. James Petersson, Elizabeth Rhoades

## Abstract

Age-related neurodegenerative disorders like Alzheimer’s disease (AD) and Parkinson’s disease (PD) are characterized by deposits of protein aggregates, or amyloid, in various regions of the brain. Traditionally, aggregation of a single protein was observed to be correlated with these different pathologies: tau in AD and α-synuclein (αS) in PD. However, there is increasing evidence that the pathologies of these two diseases overlap, and the individual proteins may promote each other’s aggregation. Both tau and αS are intrinsically disordered proteins (IDPs), lacking stable secondary and tertiary structure under physiological conditions. In this study we used a combination of biochemical and biophysical techniques to interrogate the interaction of tau with both soluble and fibrillar αS. Fluorescence correlation spectroscopy (FCS) was used to assess the interactions of specific domains of fluorescently labeled tau with full length and C-terminally truncated αS in both monomeric and fibrillar forms. We found that full-length tau as well as individual tau domains interact with monomer αS weakly, but this interaction is much more pronounced with αS seeds. This interaction does not impact tau aggregation or fibril formation. These findings provide insight into the nature of interactions between tau and αS as well as the domains responsible.

## INTRODUCTION

Alzheimer’s (AD) and Parkinson’s (PD) diseases are linked to the accumulation of aggregates of tau and α-synuclein (αS), respectively. Natively, tau is a microtubule-associated protein found predominantly in the axons of neurons that plays a critical role in microtubule stabilization and axonal transport ^1^, while αS is expressed in the presynaptic termini with a variety of putative functions, including regulation of synaptic vesicles pools ^2^, neurotransmitter release ^3^, SNARE complex assembly ^4^, and vesicle trafficking ^5^.

Aggregates of αS have been detected in disorders with tau pathology such as AD, and tau filamentation has been observed in synucleinopathies such as PD ^6 7^. This overlap in pathology translates to clinical aspects where these patients have more rapid declines as well as shortened lifespans ^8^. Studies using in vitro fibrillization monitored by K114 fluorometry followed by sedimentation analysis have also shown that these two proteins can promote each other’s aggregation, further adding to the complexity of these pathologies ^9^. However, the molecular basis for their interaction remains unknown.

Tau is organized into four major domains: the projection domain that protrudes from the microtubule surface, the proline rich region (PRR), the microtubule-binding region (MTBR) and the C-terminal domain (Fig. 1) ^10^. Within the MTBR, there are four weakly conserved repeat sequences, R1-R4. In the adult brain, alternative splicing results in six different tau isoforms, based on the absence or presence of inserts in the N-terminal region and the R2 repeat in the MTBR. Nomenclature for full-length tau is based on the inclusion of these regions. To illustrate, the longest tau isoform containing both N-terminal and all four MTBR splice regions is 2N4R. For our studies, we used 1N4R, containing all four MTBR and the N1 N-terminal insert (Fig. 1), as this is one of the most abundant isoforms in the adult brain ^11^.

**Figure 1.**
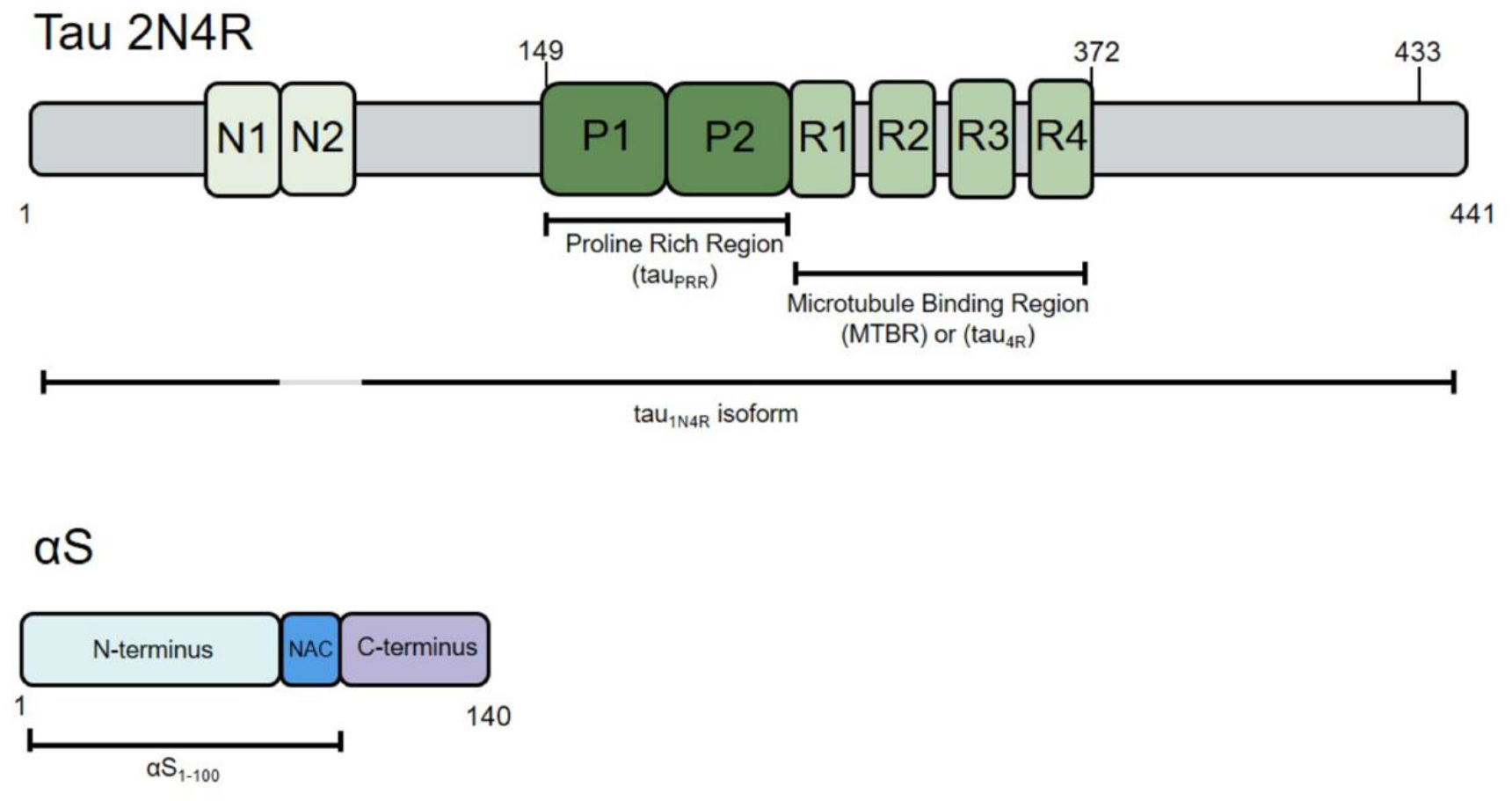
Schematic of the longest tau isoform (2N4R) and full-length αS. Shown in the tau schematic are the alternatively sliced N-terminal inserts, N1 and N2, the proline rich region (PRR) and the microtubule binding region (MTBR) with all four repeats; constructs used in this study indicated on below the schematic and all numbering of tau constructs is based on the longest isoform, 2N4R (1-441). Alexa488 fluorophore labeling is marked as tick marks above the tau schematic. On the full-length αS schematic, the major domains are labeled: the N-terminus, the NAC domain, and the C-terminus, with the αS_1-100_ construct used in this study indicated below.

αS is a 140 amino acid protein consisting of three major domains: the N-terminal domain containing imperfect repeats of the consensus sequence (KTKEGV), the hydrophobic non-amyloid-beta component or (NAC) domain, and the C-terminal domain, enriched in negative charge and proline residues^12 13^ (Fig. 1). The first ∼100 residues of αS mediate binding to lipid bilayers ^14^. In SH-SY5Y cells co-transfected with both tau and αS, an interaction was detected between the two proteins via a Bimolecular Fluorescence Complementation assay ^15^. Another study used NMR to map interactions between the C-terminus of αS and tau 1N3R as well as a tau fragment corresponding to 3R alone^16^. Further evidence for the importance of αS’s C-terminal interactions with tau comes from the observation that successive deletion of C-terminal residues reduced the amount of fibrillar αS bound to monomer tau, while the same treatment of N-terminal residues did not impact binding ^17^. This same study mapped binding of αS to tau’s central region, including PRR and MTBR ^17^.

Here, we investigate the interactions of monomer tau with monomer and fibrillar αS. To determine the roles of various domains of both proteins, we use fragments of tau consisting of the PRR (tau_PRR_), the four microtubule binding repeats (tau_4R_, also referred to as K18 in the literature^18^) and full-length 1N4R, tau_1N4R_. For αS, we use N-terminally acetylated protein, either full-length or truncated at residue 100, αS_1-100_. We also generate N-terminally acetylated αS phosphorylated at serine 129, αS pS129 (Materials and Methods). N-terminally acetylated αS is the physiological form of the protein^19^ while nearly all Lewy bodies include αS with phosphorylation at Ser129^20^.

The interaction between tau and αS is measured by fluorescence correlation spectroscopy (FCS), complimented with ensemble aggregation studies. We determine that fibrillar αS interacts much more strongly with tau than monomer αS. However, we find that αS aggregates only weakly modulate tau aggregation, suggesting that other factors may be involved in their co-aggregation in disease.

## MATERIALS AND METHODS

### Tau cloning, purification, and labeling

Tau sequences used were cloned into a lab-made pET-HT vector (a gift from L. Regan). All tau sequences contain a N-terminal His tag with a tobacco etch virus (TEV) protease cleavage site. For site specific labeling for FCS, the native cysteines at positions 291 and 322 (numbering based on full-length tau protein 2N4R [UniProt:P10636-8]) were mutated to serine and a non-native cysteine was introduced at position 372 for tau_4R_ and position 433 for tau_1N4R_. The sequence for tau_PRR_ does not contain a cysteine so one was introduced at position 149 for labeling. Growth and purification of tau constructs were based on methods described previously ^21^. A 1 L tau expression for tau_1N4R_ and 0.5 L expressions for tau_PRR_ and tau_4R_ in LB broth (Miller) supplemented with 100 µg/mL ampicillin were induced with 1 mM isopropylthio-β-galactoside (IPTG) at OD ∼0.6 for 4 hours at 37 °C. The culture was then pelleted at 4600 rpm for 20 minutes at 4 °C. The pellet was resuspended in 30 mL of Ni-NTA tau Buffer A (TBA: 50 mM Tris pH 8, 500 mM NaCl, 10 mM imidazole) with 1 mg/mL chicken egg-white lysozyme (Sigma), 1 tablet of EDTA-free protease inhibitor tablet (Roche), and 1 mM phenylmethylsulfonyl fluoride (PMSF). The resuspended pellet was sonicated on ice for 1 minute 40 seconds, 1 second on/2 seconds off, power set to 50 W. The cellular debris was removed by centrifugation at 20,000 x g for 30 minutes. The supernatant (after centrifugation) was filtered with a 0.22 µM syringe filter and added to 5-7 mL of Ni-NTA resin equilibrated with TBA, followed by incubation with rocking at 4 °C for ∼1 hour. The resin was then washed with ∼30 mL TBA and the protein was eluted with ∼15 mL Ni-NTA tau Buffer B (TBB: 50 mM Tris pH 8, 500 mM NaCl, 400 mM imidazole). The eluent was then exchanged back into TBA and concentrated to ∼1 mL using Amicon concentrators (Sigma). The His-tag was cleaved by overnight incubation at 4 °C with 1 100 μL aliquot of TEV protease (260 μM) and freshly prepared 1 mM dithiothreitol (DTT). Removal of the cleaved His-tag, as well as any uncleaved His-tagged tau, was achieved by incubation of the cleaved sample with TBA equilibrated Ni-NTA resin for 1 hour at 4 °C with rocking. The column flow through containing the cleaved tau protein was exchanged into tau Buffer C (TBC; 25 mM Tris PH 8, 100 mM NaCl) using an Amicon concentrator and concentrated down to 0.5-2 mL. The solution was filtered using a 0.22 µm filter and further purified on an ӒKTA pure FPLC system using a HiLoad Superdex 200 pg size exclusion column.

For generating labeled constructs, freshly purified tau was reduced by incubation with 1 mM DTT for 10 minutes. DTT was removed by exchanging the sample into labeling buffer (20 mM Tris pH 7.4, 50 mM NaCl, 6 M guanidine HCl) using Amicon filters. Alexa Fluor 488 maleimide in DMSO was added in five-fold molar excess to protein and incubated overnight at 4 °C with stirring. Labeled protein was exchanged into 20 mM Tris pH 7.4, 50 mM NaCl buffer, and unreacted dye was removed by passing the solution over two coupled HiTrap Desalting columns. Following purification, the samples concentration was determined using a NanoDrop™ spectrophotometer with the protein label setting Alexa 488 ((ε(494 nm) = 73000 M^-1^cm^-1^) and protein tau (ε(280 nm) = tau_1N4R_ 7450, tau_4R/PRR_ 1490 M^-1^cm^-1^) and calculated according to the Alexa Fluor ® 488 Microscale Protein Labeling Kit (A30006) using protein concentration = [A_280_-0.11(A_494_)]/A_280_) and then were aliquoted out into microcentrifuge tubes, flash frozen and stored at -80°C.

### αS cloning and purification

N-terminally acetylated αS [UniProt: P37840] was produced by co-transfection of BL21 cells with both αS and N-terminal acetyltransferase B (NatB; a gift from D. Mulvihill) complex plasmids, using ampicillin and chloramphenicol to select for colonies containing both plasmids ^22^.

The parent αS plasmid contains αS fused to a His-tagged GyrA intein from Mycobacterium xenopi (αS-intein) and was expressed and purified as previously described for non-acetylated αS ^23,24^. A 0.5 L LB broth was supplemented with 100 µg/mL ampicillin and 34 µg/mL chloramphenicol and induced with 1 mM IPTG at OD ∼0.6 overnight at 16 °C. The culture was centrifuged at 4600x g for 20 minutes at 4 °C. The pellet was resuspended in 30 mL of 40 mM Tris pH 8 supplemented with 1 tablet of EDTA-free protease inhibitor tablet (Roche), and 0.1 mM PMSF. The resuspended pellet was then sonicated on ice for 1 minute 40 seconds, 1 second on/2 seconds off, power set to 50 W. The cellular debris was removed by centrifugation at 20,000 x g for 30 minutes. The supernatant (after centrifugation) was filtered with a 0.22 µM syringe filter and added to 5-7 mL of Ni-NTA resin equilibrated with αS-intein buffer 1 (ASB1; 50 mM HEPES pH 7.5), followed by incubation with rocking at 4 °C for ∼1 hour. The resin was then washed with ∼15 mL ASB1, followed by a wash with αS-intein buffer 2 (ASB2; 50 mM HEPES pH 7.5, 5 mM imidazole). The protein was eluted with ∼12 mL Ni-NTA αS-intein buffer 3 (ASB3: 50 mM Tris pH 8, 500 mM NaCl, 400 mM imidazole). To cleave the His-tagged intein, β-mercaptoethanol (BME; 200 mM final concentration) was added to the eluent and incubated overnight (16-18 hours) at room temperature with rocking. The sample was dialyzed against 20 mM Tris pH 8.0 at 4 °C with three changes of the dialysis buffer. Removal of the cleaved His-tagged intein, as well as any uncleaved αS-intein, was achieved by incubation of the cleaved sample with Ni-NTA resin equilibrated with 20 mM Tris pH 8.0 for 1 hour at 4 °C with rocking. The column flow through (containing cleaved αS) was filtered using a 0.22 µm filter and further purified using a 5 mL HiTrap Q HP where it elutes with ∼ 300 mM NaCl.

The αS p129 was generated and purified as described above with the inclusion of a kanamycin-resistant plasmid encoding for polo-like kinase 2 (PLK2) (gift from D.T.S. Pak), a kinase that targets Ser-129^25^. Phosphorylation was confirmed via mass shift (Fig. S1) with MALDI-TOF using a Bruker rapifleX.

C-terminally truncated αS (αS_1-100_) was expressed from a T7-7 plasmid with a stop codon inserted following residue 100 ^26^. As with full-length αS, a NatB co-expression plasmid was used to generate N-terminally acetylated αS_1-100_. A culture of 0.5 L LB broth supplemented with 100 µg/mL ampicillin and 34 µg/mL chloramphenicol was induced with 320 µM IPTG at OD ∼ 0.6 and grown overnight at 16 °C. The culture was then centrifuged at 4600x g for 20 minutes at 4 °C and the resulting pellet was resuspended in 25 mL lysis buffer (20 mM Tris pH 8.0, 40 mM NaOH, 1 mM EDTA, 0.1% Triton X-100, 1mM PMSF) supplemented with 1 tablet of protease inhibitor cocktail (Roche), followed by the addition of 250 μL 1 M MgCl_2_, 250 μL 1 M CaCl_2_, and 40 µL DNase (Roche; 400U total). The sample was incubated at 37 °C for 1 hour with 250 rpm shaking. Following incubation, 500 µL of 0.5 M EDTA was added and cellular debris was removed by centrifugation at 16,900xg for 15 minutes. Ammonium sulfate cuts were used (0.116 g/mL and 0.244 g/mL) to precipitate αS in the second step. The pellet was resolubilized in αS Buffer A (ASBA; 25 mM Tris pH 8.0, 20 mM NaCl, 1mM EDTA) with 1 mM PMSF and dialyzed against ASBA overnight to remove ammonium sulfate. Dialyzed samples were then loaded onto a 5 mL HiTrap Q HP anion exchange column and eluted with αS Buffer B (ASBB; 25 mM Tris pH 8.0, 1 M NaCl, 1mM EDTA), where it elutes with ∼ 300 mM NaCl. Fractions containing αS_1-100_ were pooled and concentrated using Amicon concentrators, then loaded onto a HiLoad Superdex 200 pg size exclusion column for further final purification. Following purification, the samples were aliquoted out into microcentrifuge tubes, flash frozen and stored at -80°C.

### αS aggregation

Fibrils of αS and αS_1-100_ were generated by incubation of 100 μM protein for 7 days with agitation at 1300 rpm and 37 °C in 1x PBS (1mM KH_2_PO_4_, 3 mM Na_2_HPO_4_, 155 mM NaCl pH 7.4). Fibrils of full length αS and αS_1-100_ was imaged by EM (as described in TEM imaging below) to ensure subsequent seeds formed from fibrillar samples (Fig S2). Fibrils were stored for later use by freezing aliquots over dry ice for storage at -80 °C. Prior to use, 8 μL aliquots of resuspended fibrils were added to 392 μL of buffer (1:50 dilution, final concentration αS = 2 μM monomer units) in a microcentrifuge tube placed on ice. The diluted sample was sonicated using a QSonica microtip for 2 minutes, amplitude 50, 1 sec on /1 sec off to generated short fibers, termed seeds. For aggregation experiments, the seeds were used immediately; 50 μL of the sonicated material was mixed with 50 μL tau for a final αS seed concentration of 1 μM (monomer units).

### FCS Measurements

FCS measurements were conducted on our home-built instrument as described previously ^27^. Briefly, the power of a 488 nM DPSS laser was adjusted to ∼5 μW as measured prior to entering the microscope. Fluorescence emission was collected through the objective and separated from laser excitation using a Z488RDC long-pass dichroic and an HQ600/200M bandpass filter and focused onto the apertures of a 50 μm diameter optical fiber directly coupled to an avalanche photodiode. A digital correlator (FLEX03LQ-12, http://correlator.com) was used to generate the autocorrelation curves. All FCS measurements were carried out at ∼20 nM fluorescently labeled protein at 20°C in Nunc chambered coverslips (ThermoFisher). The chambers are pre-incubated with poly-lysine conjugated polyethylene glycol (PLL-PEG) to minimize non-specific adsorption of the protein ^28^. The buffer used for all FCS measurements consisted of 50 mM Na_2_HPO_4_, 50 mM NaCl pH 7.0, which was previously used in NMR titration experiments between αS and tau ^16^. Seeded samples were measured within 30 minutes of preparation as they appeared unstable at longer time points.

For each measurement, 25 traces of 10 s were obtained. The autocorrelation function G(*τ*) was calculated as a function of the delay time *τ* and then fit using lab-written scripts in MATLAB to a single-species diffusion equation:

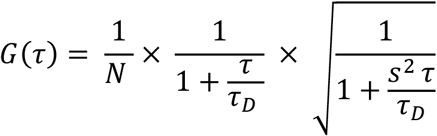

where *N* is the average number of molecules in the focal volume, *s* is the ratio of radial to axial dimensions of the focal volume, and τ_*D*_ is the translational diffusion time. At higher αS concentrations (Fig. S2) and for the seeded measurements, or even more infrequently for just labeled species only, the presence of infrequent, large bright species can disproportionally weight the averaged autocorrelation curves used in the analysis described above. Working under the premise that removal of these outliers would allow a more meaningful analysis, gathered individual diffusion times from each curve by adapting previously lab-written MATLAB codes ^27^. We then applied a non-linear regression outlier discarding algorithm known as the ROUT method on GraphPad Prism 9 to remove aberrant curves ^29^. The ROUT method combines robust regression and outlier removal to identify outliers when fitting data with nonlinear regression. This method is based on the False Discovery Rate (FDR). When there are outliers in the data, Q is the maximum desired false discovery rate. We set Q to 1%, (aiming for no more than 1% of the identified outliers to be false (are in fact just the tail of a Gaussian distribution) and at least 99% to be actual outliers (from a different distribution). The number of aberrant curves discarded for each condition is listed in Table S1.

### Aggregation Assays

Kinetic measurements of aggregation of1N4R were carried out with 5 μM tau, 100 μM DTT, and 10 μM Thioflavin T (ThT) in 1x PBS buffer, pH 7.5 in a sample volume of 100 μL. Some samples included 1.25 μM heparin from porcine intestinal mucosa (Sigma-Aldrich; average mol. wt = 18kD). Conditions with seeded αS (either full-length or αS_1-100_) included 1 μM αS seed (monomer concentration). All components were mixed in a microcentrifuge tube and then transferred to a half-area 96 well plate (Grenier) and incubated in an Ika MS3 orbital shaker set to 1300 rpm in a My Temp™ Mini digital incubator from Dot Scientific at 37 °C and monitored over the course of 72 hours. At each time point, the plate was moved to a Tecan Spark plate reader and the ThT fluorescence was measured using excitation 450 nm and emission 485 nm. After the final time point, samples were pelleted by centrifugation at maximum speed on a tabletop centrifuge (13,200 rpm) for 90 min. The supernatant was removed, and the pellet was resuspended in the original volume of buffer. Samples were supplemented with SDS to 25 mM final concentration, boiled for 20 min, and chilled on ice. All samples were analyzed by SDS-PAGE (4-12% Bis Tris, 200 V, 30 min). Gel bands were quantified with ImageJ software. A one-way ANOVA test (GraphPad Prism) was performed to test for any statistical difference in the amount of pellet in the post-aggregation gel samples.

### TEM Imaging

Following aggregation, fibril samples were pelleted in the same manner as described above, and the pellets were resuspended in 1X PBS buffer, pH 7.4. The fibrils (5 μL of 7 μM) were added to freshly glow-discharged carbon coated mesh grids and incubated for 2 minutes. Excess buffer was absorbed by wicking with filter paper. To remove excess buffer salts, grids were washed with 5 μL water, again followed by wicking. Staining of the aggregates was achieved by two rounds of incubation with 5 μL uranyl acetate (2% in water), as described above. The grids were dried under vacuum pump for 2 minutes. The post-aggregation samples were imaged on 2700x low dose and 6500x magnification and fibril samples were imaged on exposure mode and 11000x magnification on a T12 Tencai microscope.

## RESULTS

The binding of tau to both αS monomers and aggregated seeds was probed using FCS. One advantage of FCS is that allowing for the study of equilibrium protein interactions at low concentrations of fluorescent molecules, such as the nanomolar range used for the tau constructs in our study^30^. Measurements of fluorescently labeled tau_1N4R_, tau_4R_ and tau_PRR_ were made in the absence or presence of unlabeled αS monomer or αS aggregates.

### Tau interacts weakly with monomeric αS

To measure the interaction between tau and αS, the diffusion time, τ_D_, of fluorescently labeled tau was determined with increasing concentrations of monomer αS. A change in mass or hydrodynamic radius of tau upon binding αS results in a change in tau’s diffusion time, Δτ_D_ = τ_D_ (+αS) - τ_D_ (-αS). All three tau constructs bind to monomer αS, as seen by the increase in their diffusion times in its presence (Fig. 2a-c). The increase in τ_D_ is concentration dependent (Figure S3), however it is relatively weak, with high αS:tau ratios required to observe binding. We quantified this increase as the percent difference as Δτ_D_ (Table S2); at the highest concentrations of αS measured, 150 μM, increases of 7, 11 and 19% were measured for tau_1N4R_, tau_4R,_ and tau_PRR_ respectively (Table S2). To eliminate the possibility that increased viscosity due to the high concentration of αS might lead to a non-specific increase in τ_D_, enhanced green fluorescent protein (eGFP), which is not expected to interact with αS, was subjected to similar FCS measurements in the absence and presence of 150 μM αS (Fig. S4). The relative increase in τ_D_ was ∼4% (Figure S4), indicating that the larger τ_D_ increases seen for the tau constructs are likely due to at least transient interactions with αS. Interestingly, the largest relative increases in τ_D_ were observed for tau_PRR_, a domain which had not previously been identified as interacting with αS. Because of its relatively large increase, we used tau_PRR_ as a point of comparison for other αS variants. Consistent with prior studies of other tau domains, we found no change in the median diffusion time of tau_PRR_ in the presence of 150 μM αS_1-100_, underscoring the importance of the negatively charged C-terminus of αS in mediating interactions with tau_PRR_ as well. However, we were surprised to find that αS_pS129_ bound comparably to αS (Fig. S5). We had expected that the increased negative charge resulting from phosphorylation would enhance binding.

**Figure 2.**
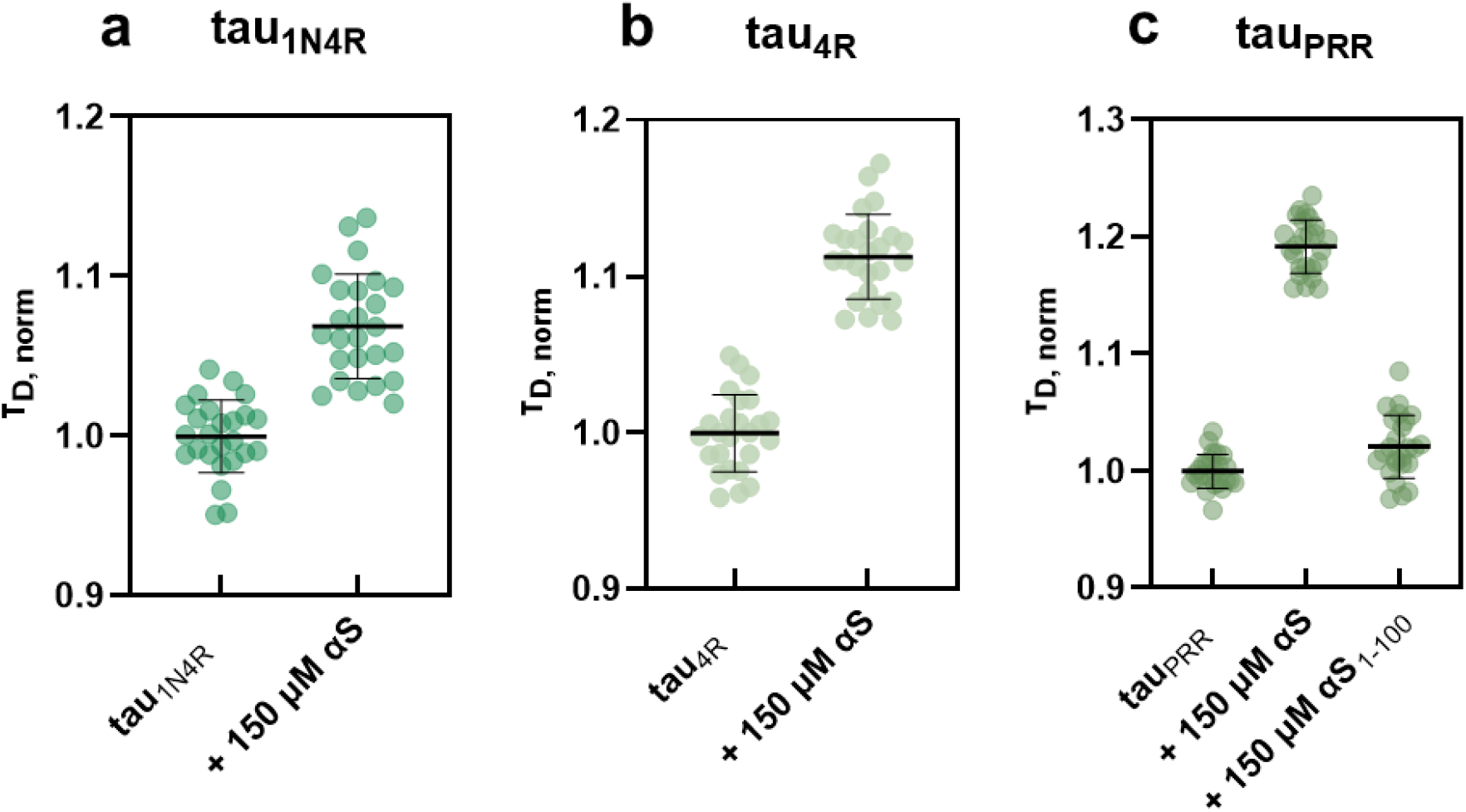
Distribution of normalized diffusion times, τ_D_, for tau_1N4R_ (a) tau_4R_ (b) and tau_PRR_ (c) in the absence and presence of 150 μM αS. For tau_PRR_, the measurement was also carried out in the presence of 150 μM αS_1-100_. Mean τ_D_ and SD calculated for a minimum of three measurements with αS. The values obtained from normalized diffusion times is found in Table S2, as well as the percent change in the presence of αS relative to tau alone.

### αS aggregates enhances interactions with tau

Aggregation of tau can be accelerated through ‘seeding’, i.e. adding tau, or even other protein aggregates to a solution of tau monomer^31^. This prompted us to investigate the interaction between tau monomers and αS aggregates. Fibrillar fragments, or seeds, of αS or αS_1-100_ were generated as described in the Materials & Methods and added to labeled tau monomer in increasing concentrations. As with the monomer αS, binding of all three tau constructs to αS seeds could be observed by an increase in τ_D_. However, in stark contrast to the monomer αS, binding could be observed even at nanomolar concentrations (monomer units) of seeds (Fig. 4, Table S5, Fig S6). As with the monomer αS constructs, binding to αS_pS129_ seeds was comparable to that of unmodified αS seeds (Fig. S7). Also consistent with the monomer protein is that deletion of the C-terminal tail of αS diminished the interaction with tau for all three constructs (Fig. 3). Control measurements with eGFP and 60 nM αS seeds resulted in only a minor increase in τ_D_, (Fig. S8). As a whole, our observations underscore the importance of αS C-terminal tail for interactions between αS and tau, and demonstrate that for fibrils, these interactions can occur at far lower concentrations than those reported in previous NMR studies ^16^.

**Figure 3.**
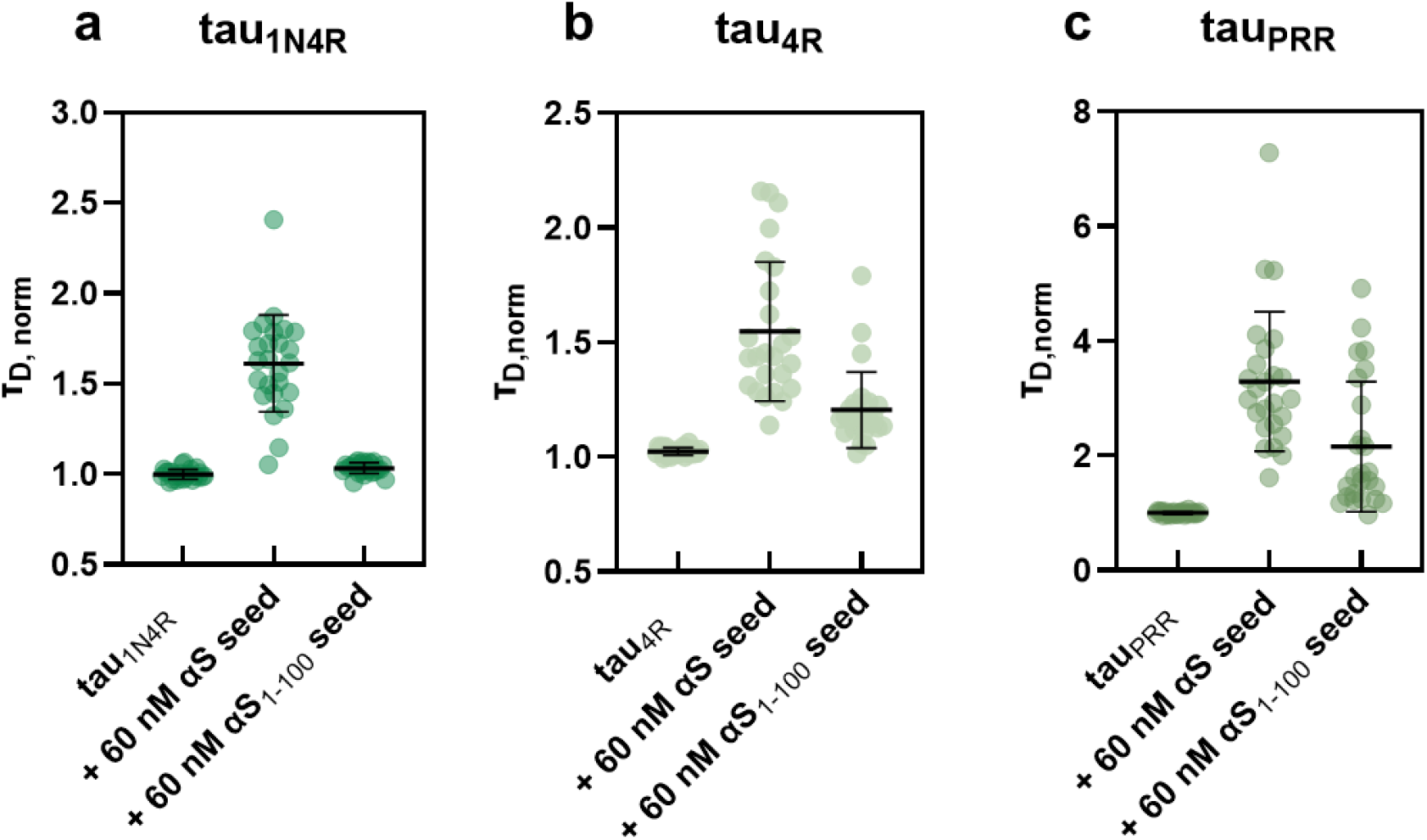
All tau constructs interact strongly with αS seeds. Individual diffusion values plotted for tau_1N4R_ a) and fragments tau_4R_ c) tau_PRR_ displaying their change in diffusion values when nanomolar unlabeled seeded αS is added, as well as C-terminally truncated seeded αS. Change in diffusion values as well as percent change compared to tau protein alone are displayed in Table S5.

**Figure 4.**
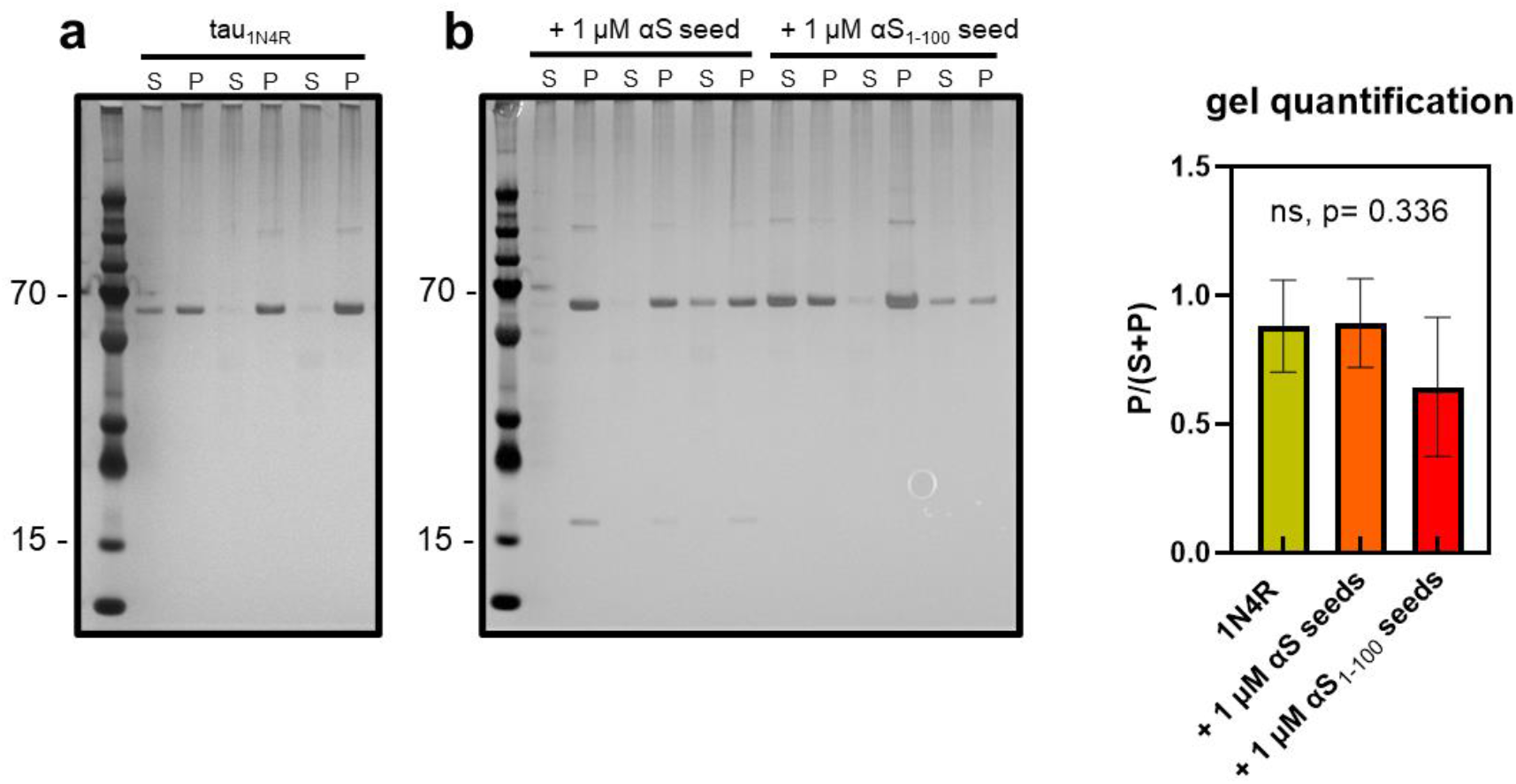
αS seeds do not impact amount of tau aggregates. SDS-PAGE gels of samples post-aggregation a) with S indicating the supernatant and P the pellet after centrifugation. The amount for each sample was quantified in triplicates using the relative amount of pellet vs. the total amount of pellet and supernatant (ImageJ) in b). 1 μM seed 1:5 molar ratio of αS to tau_1N4R_.

### αS seeds do not accelerate tau aggregation

Given the observed interaction of tau with αS seeds at nM concentrations, we sought to determine how these differences in binding affect aggregation. Aggregation was tracked using thioflavin T (ThT) fluorescence (Materials and Methods) over the course of 72 hours. Samples were pelleted following aggregation and both pellet and supernatant were visualized by SDS-PAGE. We first established conditions for aggregation of tau_1N4R_ in the absence of the commonly used molecular inducer, heparin (Fig. S9). We then compared the quantity of aggregated material in the absence and presence of seeds of αS or αS_1-100_ following 72 hours of incubation (Fig. 4). Performing a one-way ANOVA test showed there is no statistical difference in the amount of aggregated material under any of the conditions we tested (Fig. 4). With increasing molar concentration of αS seeds, we observed a slight increase in the lag time when full-length seeds were used, whereas the effect was not as pronounced when using increasing concentrations of truncated seeds in the assay (Fig S9). Additionally, a technical limitation of using ThT is its limited sensitivity to early-stage assemblies of tau. It has been shown that although tau protofibrils do bind ThT, the interaction is often weaker than observed for mature fibrils ^32^.

The pellets were also visualized by TEM following aggregation. All samples which included tau_1N4R_ alone, with heparin, with full length seeds, with truncated seeds, showed fibrillar structures (Fig. 5), further corroborating the minimal impact of the αS seeds on tau aggregation under the conditions of our assay.

**Figure 5.**
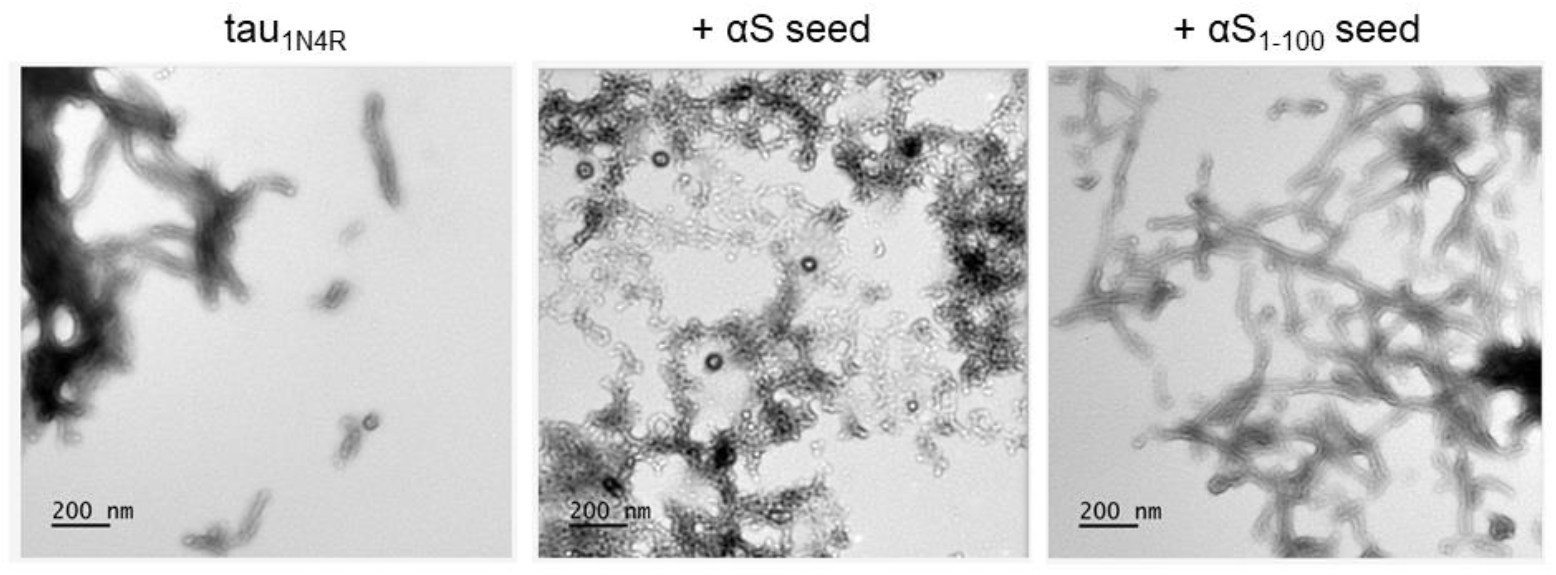
αS seeds do not impact tau fibrils formation. SDS-PAGE gels of samples post-aggregation were imaged on a T12 Tencai microscope.

## DISCUSSION

Since tau and αS were first linked to neurodegenerative disorders, both proteins have been heavily studied separately for their role in neurodegenerative diseases by a wide variety of experimental methods, but not as commonly for their roles when both are present. In this study, we have examined the interaction of tau and αS by complementing traditional aggregation with FCS. Our results have identified that 1) full-length tau, as well as individual tau domains, interact with monomeric αS weakly; 2) this interaction is much more pronounced for αS aggregates; 3) under the conditions of our assay, this interaction does not significantly impact tau aggregation or fibril formation.

We demonstrate that the PRR-MTBR of tau binds to αS (Fig 2), with the C-terminus of αS enhancing the interaction (Figs 2, 3), consistent with prior work emphasizing these regions of both proteins ^16, 17^. Our study goes on to show that the PRR and MTBR are individually capable of binding to αS. Given the uneven distribution of charged amino acids in both tau and αS (Fig S10), it is worth considering their impact on the interaction. Both the PRR and MTBR have a comparably high fraction of positively charged residues (∼0.17). However, in the PRR, there are relatively fewer negatively charged residues, resulting in it having a higher positive net charge per residue (NCPR), ∼0.14 for PRR to MTBR’s ∼0.9 (Table S2) ^33^. This may also explain why overall the PRR shows the largest shifts in diffusion times in interactions with both monomer and fibrillar αS, given the demonstrated importance of the negatively charged C-terminus of αS (NCPR ∼-0.33) (Figs 2c, 3). The somewhat surprising observation that pS129 in αS did not enhance interactions with tau (Fig S5) may also be due to the overall high negative charge within the C-terminus of αS. Specifically, the presence of the phosphate group on serine 129 only moderately increases the negative NCPR from ∼-0.33 to ∼-0.35 of this region; the high abundance of other negative charge already present within the sequence may blunt the impact of this PTM on electrostatically-driven tau interactions.

Although full-length 1N4R includes both the PRR and MTBR, it but does not show an additive enhancement in its interactions with αS (Figs 2, 3). This may be due to the presence of the net negatively charged N-terminus (NCPR∼ -0.13 for the 1N isoform; Table S2). In solution, the N-terminus of tau makes long-range electrostatic interactions with the PRR and MTBR^30^. The N-terminus thus may have repulsive interactions with the negatively charged C-terminus of αS, effectively reducing the ability of PRR and MTBR to bind to αS in the context of full-length tau.

For all tau constructs, the interaction between tau and αS was significantly enhanced for αS aggregates relative to monomer, with increases in diffusion time seen a significantly lower concentrations of aggregates (Fig 2, 3). The larger size of the aggregates certainly facilitates observation of the binding interaction, as the relative increase in τ_D_ will be larger for tau binding to αS seeds than to αS monomer. However, we observe an interaction with αS seeds at concentrations ∼1000x less than required for monomer, and we consider this observation in light of the physico-chemical properties of the aggregates. All Cryo-EM and NMR structures of αS fibers to date show the C-terminus of αS on the outside of the fiber, where it remains disordered and accessible for binding interactions^34,35^. As such, the fiber surface consists of a very high density of negatively-charged C-termini, providing multivalent binding site for positively charged biomolecules. As a consequence, it is not terribly surprising that such a surface interacts favorably with tau_PRR_ and tau_4R._ This may also provide insight into why we did not see acceleration of tau aggregation in the presence of αS seeds, namely that binding of monomer tau to fibers inhibits de novo aggregation of tau by depleting the monomer pool. FCS measurements of solutions comparable to those used for tau_1N4R_ aggregation showed a small increase in τ_D_ (Fig S11), supporting this hypothesis.

## CONCLUSIONS

Our results show that tau interacts rather weakly with monomeric αS, with a significantly enhanced interaction with αS seeds, particularly for tau_PRR_. While the MTBR is often the focus of tau studies, prior work from our lab has drawn attention to the PRR, showing that it has a critical role in binding to tubulin and tubulin polymerization^36^, as well as in binding to the aggregation inducer, polyphosphate^37^. Many of the phosphorylation modifications to tau that are used as markers of disease are found in the PRR^38^, and our work here suggests that those modifications could alter tau interactions with αS in vivo. Our results also suggest that the mechanism by which αS affects tau pathology may not necessarily be through directly seeding tau aggregation. For example, binding of tau to fibrillar αS may sequester tau away from microtubules, increasing its likelihood of it undergoing post-translational modifications that lead to aggregation. Clearly, more investigation is warranted and our own studies incorporating additional post-translational modifications and cell-based experiments are already underway.

## ASSOCIATED CONTENT

### Supporting Information

The Supporting Information is available free of charge on the ACS Publications website. Figures and tables provide protein characterization, FCS curve fits, additional diffusion data, aggregation curves, and protein charge analysis (PDF).

## ACKNOWLEDGMENT

The authors thank Ryan Kubanoff for training on the MALDI-TOF Bruker rapifleX NIH#(1S0OD030460-01). We acknowledge the use of instruments at the Electron Microscopy Resource Lab (RRID: SCR_022375).

## ABBREVIATIONS

AD: Alzheimer’s disease
PD: Parkinson’s disease
αS: α-synuclein
IDPs: intrinsically disordered proteins
FCS: Fluorescence correlation spectroscopy
PRR: proline rich region
MTBR: microtubule-binding region
NAC: non-amyloid-beta component

## REFERENCES

1. Kempf, M.; Clement, A.; Faissner, A.; Lee, G.; Brandt, R., Tau binds to the distal axon early in development of polarity in a microtubule- and microfilament-dependent manner. J Neurosci 1996, 16 (18), 5583–92.

2. Bellani, S.; Sousa, V. L.; Ronzitti, G.; Valtorta, F.; Meldolesi, J.; Chieregatti, E., The regulation of synaptic function by alpha-synuclein. Commun Integr Biol 2010, 3 (2), 106–9.

3. Cheng, F.; Vivacqua, G.; Yu, S., The role of alpha-synuclein in neurotransmission and synaptic plasticity. J Chem Neuroanat 2011, 42 (4), 242–8.

4. Burre, J.; Sharma, M.; Tsetsenis, T.; Buchman, V.; Etherton, M. R.; Sudhof, T. C., Alpha-synuclein promotes SNARE-complex assembly in vivo and in vitro. Science 2010, 329 (5999), 1663–7.

5. Scott, D.; Roy, S., alpha-Synuclein inhibits intersynaptic vesicle mobility and maintains recycling-pool homeostasis. J Neurosci 2012, 32 (30), 10129–35.

6. Arima, K.; Hirai, S.; Sunohara, N.; Aoto, K.; Izumiyama, Y.; Ueda, K.; Ikeda, K.; Kawai, M., Cellular co-localization of phosphorylated tau- and NACP/alpha-synuclein-epitopes in lewy bodies in sporadic Parkinson’s disease and in dementia with Lewy bodies. Brain Res 1999, 843 (1-2), 53–61.

7. Lippa, C. F.; Fujiwara, H.; Mann, D. M.; Giasson, B.; Baba, M.; Schmidt, M. L.; Nee, L. E.; O’Connell, B.; Pollen, D. A.; St George-Hyslop, P.; Ghetti, B.; Nochlin, D.; Bird, T. D.; Cairns, N. J.; Lee, V. M.; Iwatsubo, T.; Trojanowski, J. Q., Lewy bodies contain altered alpha-synuclein in brains of many familial Alzheimer’s disease patients with mutations in presenilin and amyloid precursor protein genes. Am J Pathol 1998, 153 (5), 1365–70.

8. Bassil, F.; Meymand, E. S.; Brown, H. J.; Xu, H.; Cox, T. O.; Pattabhiraman, S.; Maghames, C. M.; Wu, Q.; Zhang, B.; Trojanowski, J. Q.; Lee, V. M., alpha-Synuclein modulates tau spreading in mouse brains. J Exp Med 2021, 218 (1).

9. Giasson, B. I.; Forman, M. S.; Higuchi, M.; Golbe, L. I.; Graves, C. L.; Kotzbauer, P. T.; Trojanowski, J. Q.; Lee, V. M., Initiation and synergistic fibrillization of tau and alpha-synuclein. Science 2003, 300 (5619), 636–40.

10. Mandelkow, E. M.; Biernat, J.; Drewes, G.; Gustke, N.; Trinczek, B.; Mandelkow, E., Tau domains, phosphorylation, and interactions with microtubules. Neurobiol Aging 1995, 16 (3), 355–62; discussion 362-3.

11. Wheeler, J. M.; McMillan, P. J.; Hawk, M.; Iba, M.; Robinson, L.; Xu, G. J.; Dombroski, B. A.; Jeong, D.; Dichter, M. A.; Juul, H.; Loomis, E.; Raskind, M.; Leverenz, J. B.; Trojanowski, J. Q.; Lee, V. M.; Schellenberg, G. D.; Kraemer, B. C., High copy wildtype human 1N4R tau expression promotes early pathological tauopathy accompanied by cognitive deficits without progressive neurofibrillary degeneration. Acta Neuropathol Commun 2015, 3, 33.

12. Pancoe, S. X.; Wang, Y. J.; Shimogawa, M.; Perez, R. M.; Giannakoulias, S.; Petersson, E. J., Effects of Mutations and Post-Translational Modifications on alpha-Synuclein In Vitro Aggregation. J Mol Biol 2022, 434 (23), 167859.

13. Villar-Pique, A.; Lopes da Fonseca, T.; Outeiro, T. F., Structure, function and toxicity of alpha-synuclein: the Bermuda triangle in synucleinopathies. J Neurochem 2016, 139 Suppl 1, 240–255.

14. Rhoades, E.; Ramlall, T. F.; Webb, W. W.; Eliezer, D., Quantification of alpha-synuclein binding to lipid vesicles using fluorescence correlation spectroscopy. Biophys J 2006, 90 (12), 4692–700.

15. Torres-Garcia, L.; Jm, P. D.; Brandi, E.; Haikal, C.; Mudannayake, J. M.; Bras, I. C.; Gerhardt, E.; Li, W.; Svanbergsson, A.; Outeiro, T. F.; Gouras, G. K.; Li, J. Y., Monitoring the interactions between alpha-synuclein and Tau in vitro and in vivo using bimolecular fluorescence complementation. Sci Rep 2022, 12 (1), 2987.

16. Lu, J.; Zhang, S.; Ma, X.; Jia, C.; Liu, Z.; Huang, C.; Liu, C.; Li, D., Structural basis of the interplay between alpha-synuclein and Tau in regulating pathological amyloid aggregation. J Biol Chem 2020, 295 (21), 7470–7480.

17. Oikawa, T.; Nonaka, T.; Terada, M.; Tamaoka, A.; Hisanaga, S.; Hasegawa, M., alpha-Synuclein Fibrils Exhibit Gain of Toxic Function, Promoting Tau Aggregation and Inhibiting Microtubule Assembly. J Biol Chem 2016, 291 (29), 15046–56.

18. Luo, Y.; Ma, B.; Nussinov, R.; Wei, G., Structural Insight into Tau Protein’s Paradox of Intrinsically Disordered Behavior, Self-Acetylation Activity, and Aggregation. J Phys Chem Lett 2014, 5 (17), 3026–3031.

19. Iyer, A.; Sidhu, A.; Subramaniam, V., How important is the N-terminal acetylation of alpha-synuclein for its function and aggregation into amyloids? Front Neurosci 2022, 16, 1003997.

20. Anderson, J. P.; Walker, D. E.; Goldstein, J. M.; de Laat, R.; Banducci, K.; Caccavello, R. J.; Barbour, R.; Huang, J.; Kling, K.; Lee, M.; Diep, L.; Keim, P. S.; Shen, X.; Chataway, T.; Schlossmacher, M. G.; Seubert, P.; Schenk, D.; Sinha, S.; Gai, W. P.; Chilcote, T. J., Phosphorylation of Ser-129 is the dominant pathological modification of alpha-synuclein in familial and sporadic Lewy body disease. J Biol Chem 2006, 281 (40), 29739–52.

21. Melo, A. M.; Coraor, J.; Alpha-Cobb, G.; Elbaum-Garfinkle, S.; Nath, A.; Rhoades, E., A functional role for intrinsic disorder in the tau-tubulin complex. Proc Natl Acad Sci U S A 2016, 113 (50), 14336–14341.

22. Trexler, A. J.; Rhoades, E., N-Terminal acetylation is critical for forming alpha-helical oligomer of alpha-synuclein. Protein Sci 2012, 21 (5), 601–5.

23. Batjargal, S.; Walters, C. R.; Petersson, E. J., Inteins as traceless purification tags for unnatural amino acid proteins. J Am Chem Soc 2015, 137 (5), 1734–7.

24. Haney, C. M.; Wissner, R. F.; Warner, J. B.; Wang, Y. J.; Ferrie, J. J.; d, J. C.; Karpowicz, R. J.; Lee, V. M.; Petersson, E. J., Comparison of strategies for non-perturbing labeling of alpha-synuclein to study amyloidogenesis. Org Biomol Chem 2016, 14 (5), 1584–92.

25. Samuel, F.; Flavin, W. P.; Iqbal, S.; Pacelli, C.; Sri Renganathan, S. D.; Trudeau, L. E.; Campbell, E. M.; Fraser, P. E.; Tandon, A., Effects of Serine 129 Phosphorylation on alpha-Synuclein Aggregation, Membrane Association, and Internalization. J Biol Chem 2016, 291 (9), 4374–85.

26. Trexler, A. J.; Rhoades, E., Alpha-synuclein binds large unilamellar vesicles as an extended helix. Biochemistry 2009, 48 (11), 2304–6.

27. Li, X. H.; Rhoades, E., Heterogeneous Tau-Tubulin Complexes Accelerate Microtubule Polymerization. Biophys J 2017, 112 (12), 2567–2574.

28. Wickramasinghe, S. P.; Rhoades, E., Measuring Interactions Between Tau and Aggregation Inducers with Single-Molecule Forster Resonance Energy Transfer. Methods Mol Biol 2020, 2141, 755–775.

29. Motulsky, H. J.; Brown, R. E., Detecting outliers when fitting data with nonlinear regression - a new method based on robust nonlinear regression and the false discovery rate. BMC Bioinformatics 2006, 7, 123.

30. Elbaum-Garfinkle, S.; Rhoades, E., Identification of an aggregation-prone structure of tau. J Am Chem Soc 2012, 134 (40), 16607–13.

31. Mirbaha, H.; Chen, D.; Morazova, O. A.; Ruff, K. M.; Sharma, A. M.; Liu, X.; Goodarzi, M.; Pappu, R. V.; Colby, D. W.; Mirzaei, H.; Joachimiak, L. A.; Diamond, M. I., Inert and seed-competent tau monomers suggest structural origins of aggregation. Elife 2018, 7.

32. Gade Malmos, K.; Blancas-Mejia, L. M.; Weber, B.; Buchner, J.; Ramirez-Alvarado, M.; Naiki, H.; Otzen, D., ThT 101: a primer on the use of thioflavin T to investigate amyloid formation. Amyloid 2017, 24 (1), 1–16.

33. Holehouse, A. S.; Das, R. K.; Ahad, J. N.; Richardson, M. O.; Pappu, R. V., CIDER: Resources to Analyze Sequence-Ensemble Relationships of Intrinsically Disordered Proteins. Biophys J 2017, 112 (1), 16–21.

34. Tuttle, M. D.; Comellas, G.; Nieuwkoop, A. J.; Covell, D. J.; Berthold, D. A.; Kloepper, K. D.; Courtney, J. M.; Kim, J. K.; Barclay, A. M.; Kendall, A.; Wan, W.; Stubbs, G.; Schwieters, C. D.; Lee, V. M.; George, J. M.; Rienstra, C. M., Solid-state NMR structure of a pathogenic fibril of full-length human alpha-synuclein. Nat Struct Mol Biol 2016, 23 (5), 409–15.

35. Li, B.; Ge, P.; Murray, K. A.; Sheth, P.; Zhang, M.; Nair, G.; Sawaya, M. R.; Shin, W. S.; Boyer, D. R.; Ye, S.; Eisenberg, D. S.; Zhou, Z. H.; Jiang, L., Cryo-EM of full-length alpha-synuclein reveals fibril polymorphs with a common structural kernel. Nat Commun 2018, 9 (1), 3609.

36. McKibben, K. M.; Rhoades, E., Independent tubulin binding and polymerization by the proline-rich region of Tau is regulated by Tau’s N-terminal domain. J Biol Chem 2019, 294 (50), 19381–19394.

37. Wickramasinghe, S. P.; Lempart, J.; Merens, H. E.; Murphy, J.; Huettemann, P.; Jakob, U.; Rhoades, E., Polyphosphate Initiates Tau Aggregation through Intra- and Intermolecular Scaffolding. Biophys J 2019, 117 (4), 717–728.

38. Wesseling, H.; Mair, W.; Kumar, M.; Schlaffner, C. N.; Tang, S.; Beerepoot, P.; Fatou, B.; Guise, A. J.; Cheng, L.; Takeda, S.; Muntel, J.; Rotunno, M. S.; Dujardin, S.; Davies, P.; Kosik, K. S.; Miller, B. L.; Berretta, S.; Hedreen, J. C.; Grinberg, L. T.; Seeley, W. W.; Hyman, B. T.; Steen, H.; Steen, J. A., Tau PTM Profiles Identify Patient Heterogeneity and Stages of Alzheimer’s Disease. Cell 2020, 183 (6), 1699–1713 e13.

