## Supplementary Information for "Nanomolar interactions of alpha-synuclein fibrils to tau determined by FCS"

**Figure S1.** MALDI-TOF spectra of phosphorylation of αS pS129……………………………S3

**Figure S2.** Representative individual autocorrelation curves with monomeric αS………………S4

**Table S1.** The total number of FCS curves collected per condition……………………………..S5

**Figure S3.** $\tau_D$ for monomer $\text{tau}_{4R}$ and $\text{tau}_{PRR}$ at increasing concentrations………..……………S6

**Table S2.** $\tau_D$ of $\text{tau}_{1N4R}$, $\text{tau}_{4R}$, and $\text{tau}_{PRR}$ with and without 150 μM αS………………………...S7

**Table S3.** $\tau_D$ of $\text{tau}_{PRR}$ and $\text{tau}_{4R}$ at increasing concentrations of αS……………………….…S8

**Figure S4.** No interaction of eGFP with monomeric αS………………………………………...S9

**Figure S5.** $\tau_D$ with monomeric αS pS129…………………………………….……...……S10

**Table S4.** Salt test in the buffer…………………………………………………………………S11

**Table S5.** $\tau_D$ of $\text{tau}_{1N4R}$, $\text{tau}_{4R}$, and $\text{tau}_{PRR}$ with and without 60 nM seeded αS…………………S12

**Figure S6.** $\tau_D$ of $\text{tau}_{1N4R}$, $\text{tau}_{4R}$, $\text{tau}_{PRR}$ at increasing αS seed concentrations…………………..S13

**Figure S7.** $\tau_D$ of $\text{tau}_{PRR}$ with seeded αS pS129………………………………..……………...S14

**Figure S8.** $\tau_D$ of eGFP with seeded αS added…………………………………..……………S15

**Figure S9.** Seeded aS/tau aggregation plots at increasing molar ratios…………………..…..S16

**Fig S10**. Charge plots of αS and tau generated using CIDER[34]…………………………………S17

**Figure S11.** $\tau_D$ using aggregation conditions……………………………………….……..…S18

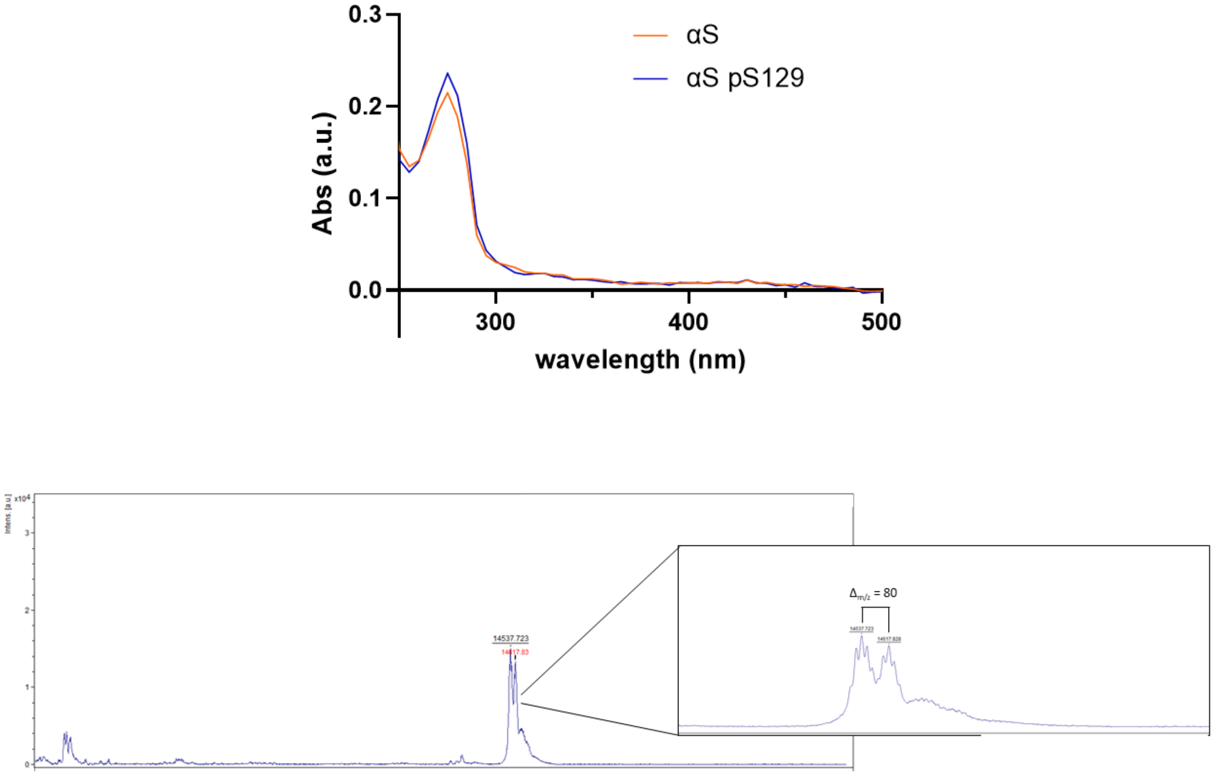


**Figure S1.** Absorbance spectra of WT acetylated αS and pS129 used to calculated concentrations of both proteins prior to MALDI-TOF (ε(280 nm) = 5960 $M^{-1}cm^{-1}$). Both proteins were mixed in equimolar amounts and spotted for MADLI-TOF analysis. The relatively equal spectral peak heights of the MALDI-TOF spectra and the expected mass shift indicate that most, if not all, the αS is phosphorylated after the co-expression of NatB and PLK2 plasmids (see Methods for details).

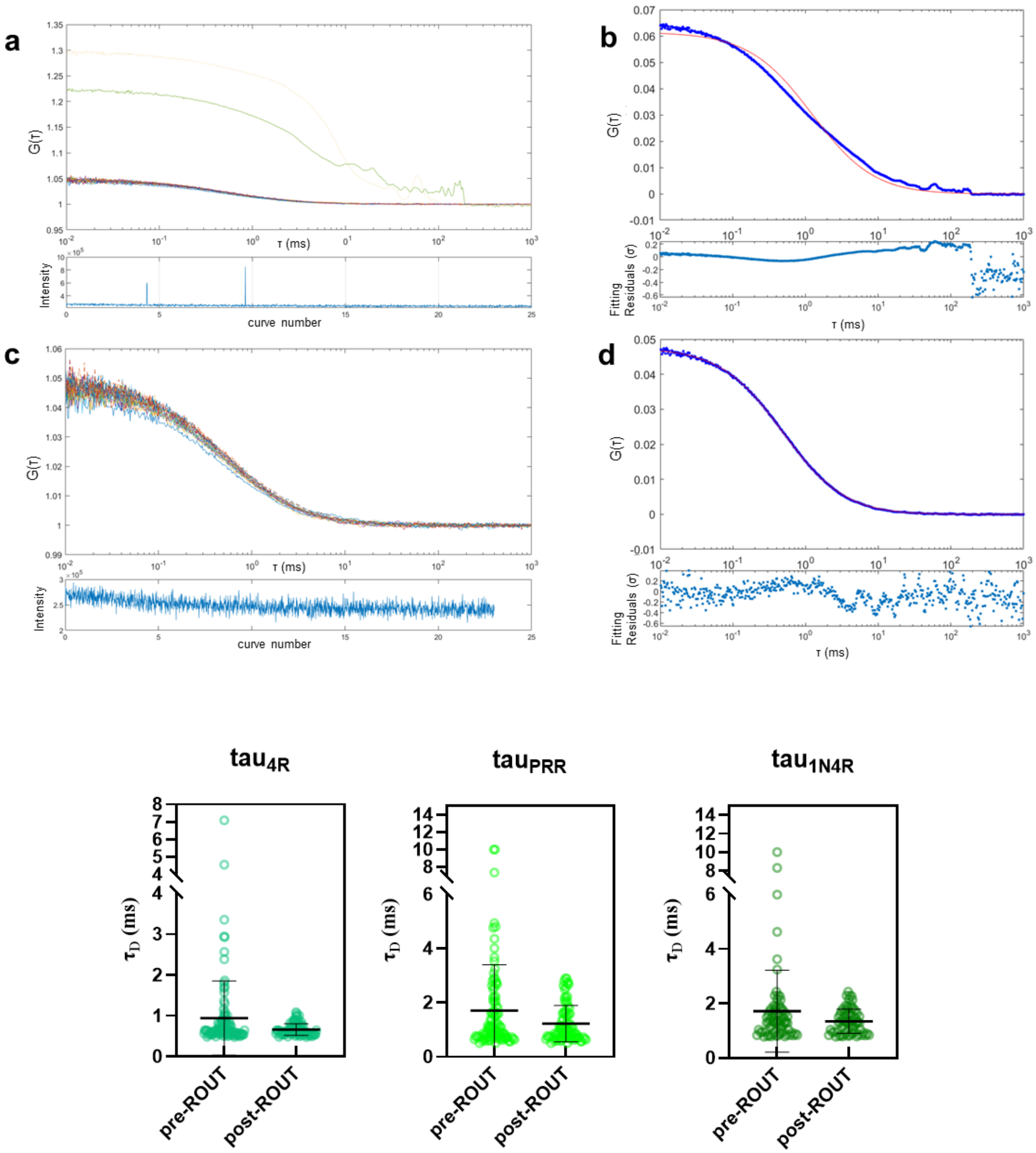


**Figure S2.** Aberrant curves disproportionally weigh the averaged autocorrelation at higher synuclein concentrations. Representative individual autocorrelation curves using labeled tau$_{4R}$ with 100 µM αS added a), averaged correlation curve in blue b) before ROUT outlier testing is performed. Prior to discarding curves, the averaged curve (blue) was not well fit with a single component 3D diffusion model (red). Post ROUT individual curves are shown in c) and averaged (blue) and fit (red) post ROUT in d). In the graphs below, representative examples of individual diffusion values along with the mean and the standard distribution obtained for each tau construct pre and post ROUT outlier testing when 60 nM αS seed is added. The majority of the data points fall within the main distribution, so that discarding aberrant curves with extremely high $\tau_D$ values does not change interpretation of the data.

| construct | Total curves collected | Total curves discarded | construct | Total curves collected | Total curves discarded | construct | Total curves collected | Total curves discarded |
|---|---|---|---|---|---|---|---|---|
| tau$_{PRR}$ | 725 | 4 | tau$_{4R}$ | 275 | 4 | tau$_{1N4R}$ | 275 | 6 |
| + 50 μM αS | 150 | 1 | + 50 μM αS | 75 | 0 | + 150 μM αS | 75 | 0 |
| + 50 μM αS$_{pS129}$ | 75 | 1 | + 100 μM αS | 75 | 2 | + 20 nM αS seed | 75 | 4 |
| + 100 μM αS | 150 | 0 | + 150 μM αS | 75 | 0 | + 20 nM αS$_{1\text{-}100}$ seed | 75 | 1 |
| + 100 μM αS$_{pS129}$ | 75 | 0 | + 20 nM αS seed | 75 | 4 | + 60 nM αS seed | 75 | 7 |
| + 150 μM αS | 150 | 1 | + 20 nM αS$_{1\text{-}100}$ seed | 75 | 5 | + 60 nM αS$_{1\text{-}100}$ seed | 75 | 5 |
| + 150 μM αS$_{1\text{-}100}$ | 75 | 0 | + 60 nM αS seed | 100 | 14 | + 100 nM αS seed | 75 | 9 |
| + 150 μM αS$_{pS129}$ | 75 | 0 | + 60 nM αS$_{1\text{-}100}$ seed | 100 | 11 | + 100 nM αS$_{1\text{-}100}$ seed | 75 | 16 |
| + 20 nM αS seed | 150 | 10 | + 100 nM αS seed | 75 | 9 | + 1 μM αS, 4.98 μM tau$_{1N4R}$ | 75 | 2 |
| + 20 nM αS$_{pS129}$ seed | 75 | 8 | + 100 nM αS$_{1\text{-}100}$ seed | 75 | 7 | + 1 μM αS$_{1\text{-}100}$, 4.98 μM tau$_{1N4R}$ | 75 | 1 |
| + 20 nM αS$_{1\text{-}100}$ seed | 75 | 5 | eGFP alone | 150 | 0 | | | |
| + 60 nM αS seed | 200 | 17 | + 150 μM αS | 75 | 0 | | | |
| + 60 nM αS$_{pS129}$ seed | 100 | 10 | + 60 nM αS seed | 75 | 0 | | | |
| + 60 nM αS$_{1\text{-}100}$ seed | 100 | 9 | | | | | | |
| + 100 nM αS seed | 150 | 18 | | | | | | |
| + 100 nM αS$_{pS129}$ seed | 75 | 11 | | | | | | |
| + 100 nM αS$_{1\text{-}100}$ seed | 75 | 12 | | | | | | |

**Table S1.** Analysis of autocorrelation curves discarded through ROUT analysis. The table contains the total number of curves collected per condition, as well as the number of curves discarded using the ROUT method.

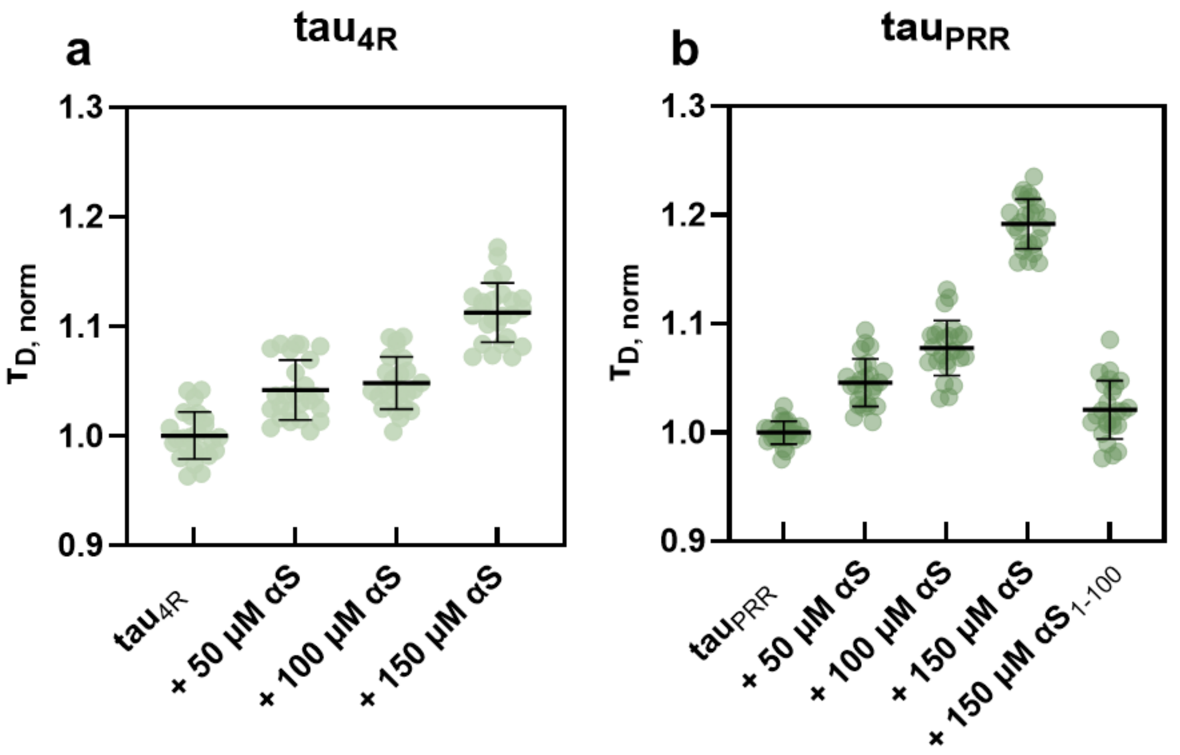


**Figure S3.** Tau binds weakly to αS monomer in a concentration dependent manner. Unlabeled αS was added to fluorescently labeled tau as described in the Materials & Methods. Normalized $\tau_D$ for a) $tau_{4R}$ and b) $tau_{PRR}$.

| Tau construct | - αS $\tau_{D, norm}$ | + αS $\tau_{D, norm}$ | + αS % diff | + $\alpha S_{1-100}$ $\tau_{D, norm}$ | + $\alpha S_{1-100}$ % diff |
|---|---|---|---|---|---|
| $tau_{1N4R}$ | 1.00±0.041 | 1.07±0.06 | 7 | - | - |
| $tau_{4R}$ | 1.00±0.038 | 1.11±0.054 | 11 | - | - |
| $tau_{PRR}$ | 1.00±0.035 | 1.19±0.086 | 19 | 1.02±0.044 | 2 |

**Table S2.** Quantification of binding at 150 μM αS monomer. Mean $\tau_{D, norm}$ and SD calculated for a minimum of three measurements with increasing concentration of αS. % diff = [$\tau_D$(+αS)-$\tau_D$(-αS)]/$\tau_D$(-αS). $Tau_{PRR}$ was also measured with 150 μM $\alpha S_{1-100}$.

| construct | $\tau_{D,\ norm}$ | % diff |
|---|---|---|
| $tau_{4R}$ | 1.00±0.038 | - |
| + 50 μM αS | 1.04±0.044 | 4 |
| + 100 μM αS | 1.05±0.043 | 5 |
| + 150 μM αS | 1.11±0.054 | 11 |
| $tau_{PRR}$ | 1.00±0.035 | - |
| + 50 μM αS | 1.05±0.047 | 5 |
| + 100 μM αS | 1.08±0.048 | 8 |
| + 150 μM αS | 1.19±0.086 | 19 |
| + 150 μM $\alpha S_{1\text{-}100}$ | 1.02±0.044 | 2 |

**Table S3.** Quantification of binding $tau_{4R}$ and $tau_{PRR}$ with increasing concentrations of αS monomer. Mean $\tau_{D,\ norm}$ and SD calculated for a minimum of three measurements for each concentration αS. %diff is calculated as $[\tau_D(+\alpha S)-\tau_D(-\alpha S)]/\tau_D(-\alpha S)$

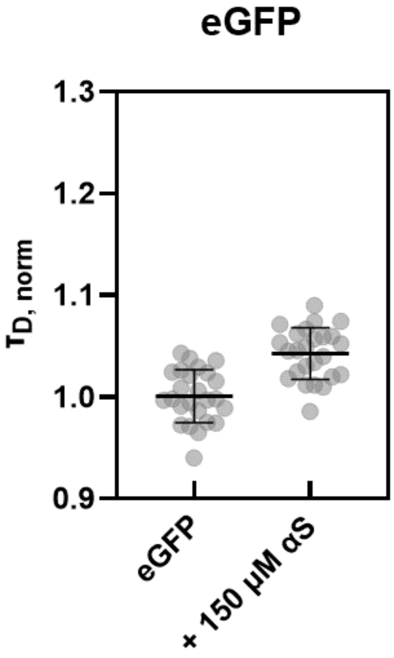


| construct | $\tau_{D,\,norm}$ - αS | $\tau_{D,\,norm}$ + αS | % diff + αS |
|---|---|---|---|
| eGFP | 1.00±0.052 | 1.04±0.053 | 4 |

**Figure S4.** eGFP has minimal interactions with αS monomer. Unlabeled αS was added to eGFP. Plots shown $\tau_{D,\,norm}$ for a) eGFP and b) eGFP with 150 μM αS. Mean $\tau_{D,\,norm}$ and SD calculated for a minimum of three measurements with αS. %diff = [$\tau_D$(+αS)-$\tau_D$(-αS)]/$\tau_D$(-αS).

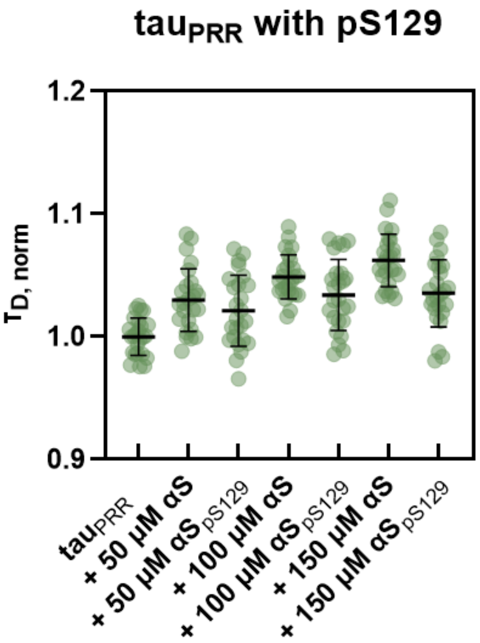


| construct | $\tau_{D,\ norm}$ | % diff |
|---|---|---|
| tau$_{PRR}$ | 1.00±0.04 | - |
| + 50 μM αS | 1.03±0.044 | 3 |
| + 100 μM αS | 1.05±0.039 | 5 |
| + 150 μM αS | 1.06±0.047 | 6 |
| + 50 μM αS$_{pS129}$ | 1.02±0.049 | 2 |
| + 100 μM αS$_{pS129}$ | 1.03±0.045 | 3 |
| + 150 μM αS$_{pS129}$ | 1.04±0.041 | 4 |

**Figure S5.** Phosphorylation of monomer αS at S129 does not enhance interactions with tau$_{PRR}$. Unlabeled αS was added at concentrations indicated on plot. Both αS and αS$_{pS129}$ show mild, concentration-dependent increases in binding, reflecting weak interactions. Mean $\tau_{D,\ norm}$ and SD calculated for a minimum of three measurements with increasing concentration of αS. %diff= $[\tau_D(+\alpha S)-\tau_D(-\alpha S)]/\tau_D(-\alpha S)$.

| construct | $\tau_{D, norm}$ | % diff |
|---|---|---|
| $tau_{PRR}$ | 0.433 | - |
| + 150 μM αS | 0.456 | 5.3 |
| + 150 μM αS with extra salt removal | 0.483 | 11.5 |

**Table S4.** Binding of $tau_{PRR}$ to αS is salt-sensitive. We note a lower overall $\tau_D$ for the measurements shown in Fig. S5 and suspected that buffer was not being as effectively exchanged by new centrifugal devices. We tested this by deliberately altering the salt in the buffer in the first condition then removed the extra salt with the new centrifugal devices. These data confirmed our hypothesis, drawing attention to the role of electrostatics in interactions between tau and αS. $\%diff=[\tau_D(+\alpha S)-\tau_D(-\alpha S)]/\tau_D(-\alpha S)$.

| Tau construct + 60 nM seed | $\tau_{D,\ norm}$ - αS seed | $\tau_{D,\ norm}$ + αS seed | $\tau_{D,\ norm}$ + $\alpha S_{1\text{-}100}$ seed | % diff + αS seed | % diff + $\alpha S_{1\text{-}100}$ seed |
|---|---|---|---|---|---|
| $tau_{1N4R}$ | 1.00±0.044 | 1.63±0.559 | 1.04±0.084 | 63 | 4 |
| $tau_{4R}$ | 1.02±0.055 | 1.54±0.628 | 1.22±0.368 | 52 | 20 |
| $tau_{PRR}$ | 1.01±0.049 | 3.25±2.28 | 2.13±2.55 | 222 | 111 |

**Table S5.** Quantification of binding at 60 nM αS seeds. Unlabeled αS seeds, both full-length and $\alpha S_{1\text{-}100}$, were added to labeled tau solutions. Mean $\tau_{D,\ norm}$ and SD calculated for a minimum of three measurements with increasing concentration of αS. % diff = $[\tau_D(+\alpha S)-\tau_D(-\alpha S)]/\tau_D(-\alpha S)$ .

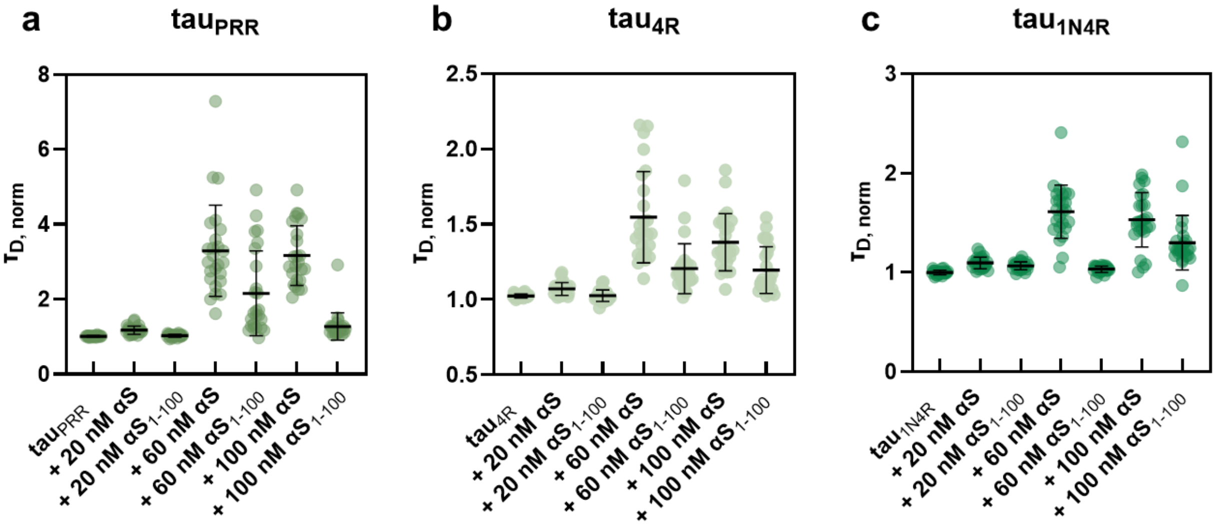


| Tau + seed conc. | $\tau_{D,\ norm}$ + αS | $\tau_{D,\ norm}$ + $\alpha S_{1\text{-}100}$ | % diff + αS | % diff + $\alpha S_{1\text{-}100}$ |
|---|---|---|---|---|
| $tau_{1N4R}$ (1.00±0.044) | - | - | - | - |
| 20 nM | 1.09±0.147 | 1.07±0.111 | 9 | 7 |
| 60 nM | 1.63±0.559 | 1.04±0.084 | 63 | 4 |
| 100 nM | 1.54±0.63 | 1.30±0.429 | 54 | 30 |
| $tau_{4R}$ (1.02±0.055) | - | - | - | - |
| 20 nM | 1.07±0.078 | 1.03±0.053 | 5 | 1 |
| 60 nM | 1.54±0.628 | 1.22±0.368 | 52 | 20 |
| 100 nM | 1.37±0.361 | 1.19±0.306 | 35 | 17 |
| $tau_{PRR}$ (1.01±0.049) | - | - | - | - |
| 20 nM | 1.18±0.261 | 1.03±0.068 | 17 | 2 |
| 60 nM | 3.25±2.28 | 2.13±2.55 | 222 | 111 |
| 100 nM | 3.18±1.48 | 1.24±0.356 | 215 | 23 |

**Figure S6.** Tau binds to αS seeds. Unlabeled αS seeds were added to labeled tau solutions. Individual $\tau_{D,\ norm}$ values are plotted for a) $tau_{1N4R}$ b) $tau_{4R}$ and c) $tau_{PRR}$ at αS seed (full-length and $\alpha S_{1\text{-}100}$) concentrations indicated on plot. .Mean $\tau_{D,\ norm}$ and SD calculated for a minimum of three measurements with increasing concentration of αS. %diff=τ[$\tau_D$(+αS)-$\tau_D$(-αS)]/$\tau_D$(-αS).

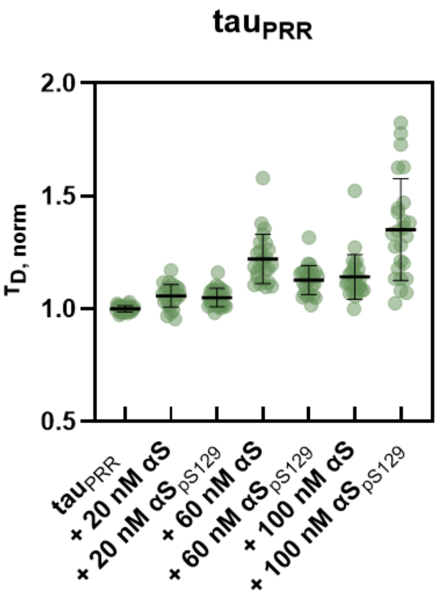


| $tau_{PRR}$ + seed conc. | $\tau_{D,\ norm}$ + αS | $\tau_{D,\ norm}$ + $αS_{pS129}$ | % diff + αS | % diff + $αS_{pS129}$ |
|---|---|---|---|---|
| $tau_{PRR}$ (1.00±0.04) | - | - | - | - |
| 20 nM | 1.06±0.079 | 1.05±0.069 | 6 | 5 |
| 60 nM | 1.22±0.235 | 1.13±0.114 | 22 | 13 |
| 100 nM | 1.13±0.12 | 1.35±0.334 | 13 | 35 |

**Figure S7.** Quantification of $tau_{PRR}$ binding to $αS_{pS129}$ seeds. Plots of $\tau_{D,\ norm}$ for $tau_{PRR}$ at αS (unmodified and $αS_{pS129}$) seed concentrations of 20, 60, and 100 nM. Mean $\tau_{D,norm}$ and SD calculated for a minimum of three measurements with increasing concentration of αS. %diff $=\tau_D(+αS) - \tau_D(-αS)\ ]/\tau_D(-αS)$. Note the lower overall $\tau_{D,\ norm}$ as seen with monomer pS129 (Figure S5); the overall trend of observed with the monomer protein is seen with seeds as well.

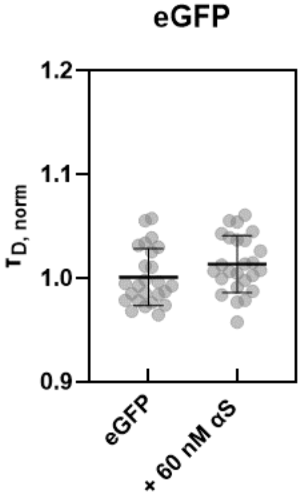


| construct | $\tau_{D, norm}$ - αS seed | $\tau_{D, norm}$ + αS seed | % diff + αS seed |
|---|---|---|---|
| eGFP | 1.00±0.048 | 1.01±0.056 | 1 |

**Figure S8.** eGFP does not bind to αS seeds. Unlabeled αS seeds were added to eGFP. Mean $\tau_{D, norm}$ and SD calculated for a minimum of three measurements with αS. %diff=[$\tau_D$(+αS)-$\tau_D$(αS)]/$\tau_D$(-αS)

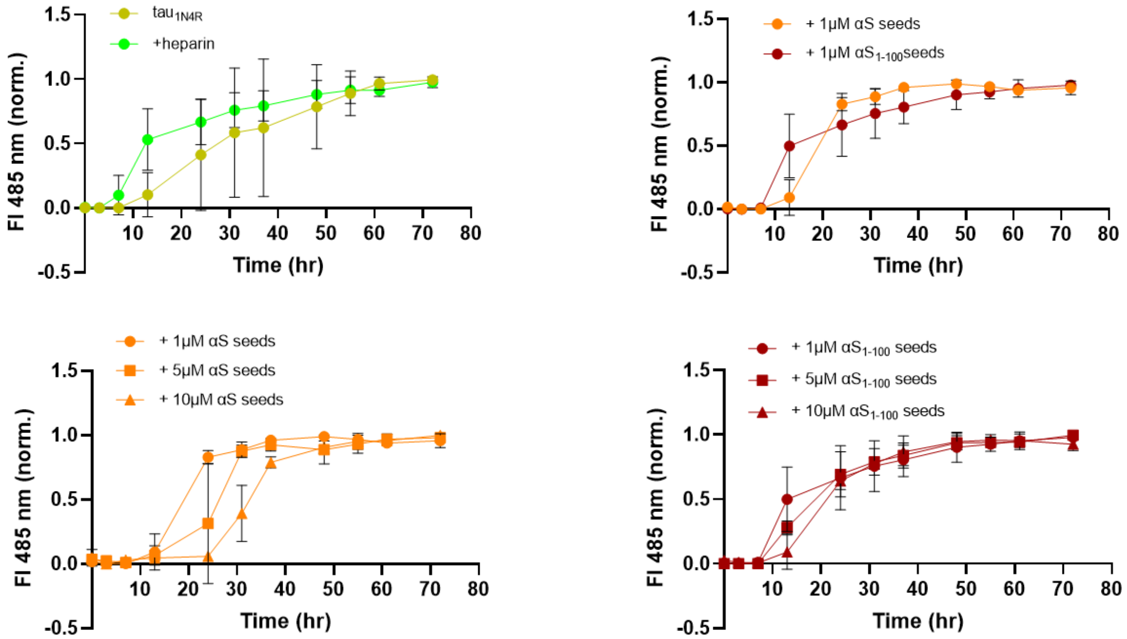


**Figure S9.** αS seeds do not significantly impact kinetics of tau aggregation. Time-points from of tau aggregation measurements using 5 μM tau$_{1N4R}$ (olive) and 5 μM tau$_{1N4R}$ (olive) with 1.25 μM heparin (green) tracked using Thioflavin T fluorescence. For experiments with αS seeds, heparin was not included in the solutions. Seeded reactions all used 5 μM tau$_{1N4R}$ and various amounts of αS seeds, as indicated on the plots above. Measurements were carried out in triplicates.

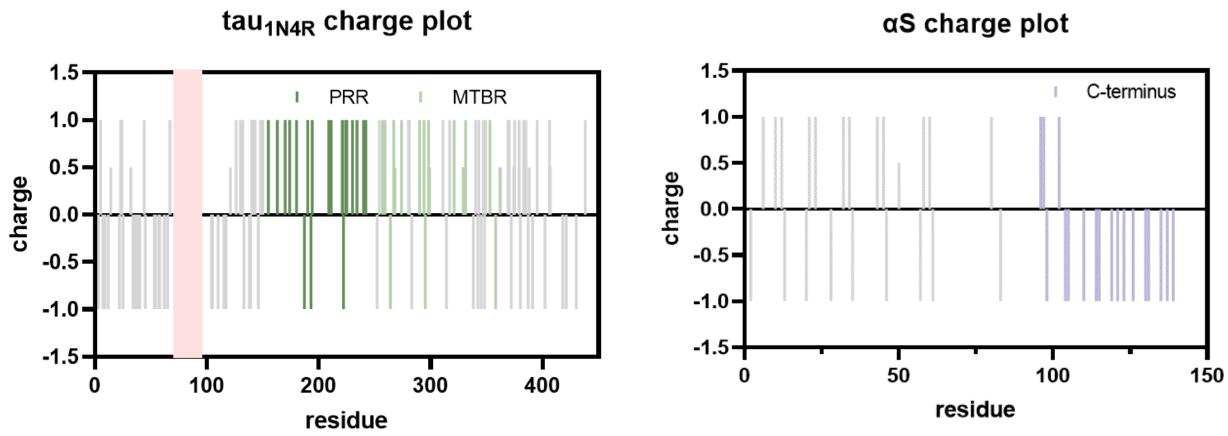


| construct | length | NCPR | κ | f+ | f- |
|---|---|---|---|---|---|
| $tau_{1N4R}$ | 414 | 0.014 | 0.181 | 0.138 | 0.123 |
| $tau_{4R}$ | 133 | 0.075 | 0.107 | 0.158 | 0.083 |
| $tau_{PRR}$ | 98 | 0.143 | 0.111 | 0.173 | 0.030 |
| αS | 140 | -0.064 | 0.172 | 0.107 | 0.171 |
| $\alpha S_{1\text{-}100}$ | 100 | 0.040 | 0.082 | 0.140 | 0.100 |
| $\alpha S_{101\text{-}140}$ | 40 | -0.325 | 0.091 | 0.025 | 0.350 |

**Fig S10**. Charge plots generated using CIDER[34] outline the distribution of positive versus negative charges in the PRR of tau compared to the repeats, as well as the distribution in the domains of αS. The numbering for tau is for the longest tau isoform, 2N4R; 1N4R lacks one of the N-terminal inserts (indicated in pink on the plot). CIDER analysis for tau and αS constructs are listed in the table, NCPR (net charge per residue), κ (kappa), f+ (fraction of positive residues), and f- (fraction of negative residues).

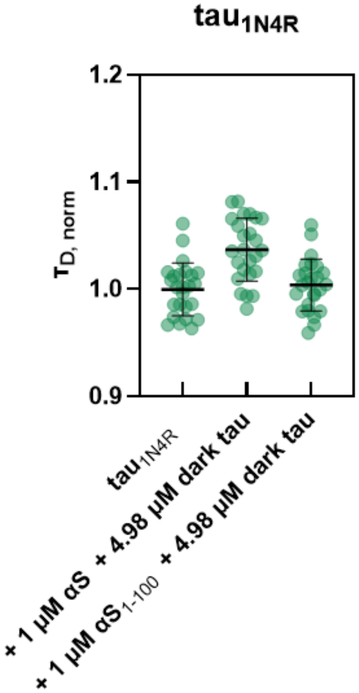


| construct | $\tau_{D,\ norm}$ | % diff |
|---|---|---|
| $tau_{1N4R}$ | 1.00±0.044 | - |
| + 1 μM αS, 4.98 μM $tau_{1N4R}$ | 1.04±0.055 | 4 |
| + 1 μM $\alpha S_{1-100}$, 4.98 μM $tau_{1N4R}$ | 1.00±0.054 | 0 |

**Figure S11.** $Tau_{1N4R}$ binds to αS seeds under aggregation conditions. FCS measurements were made under conditions comparable to those used for aggregation. Unlabeled tau allowed for high total tau concentrations, while maintaining low fluorescence levels. Plots of individual normalized $\tau_D$ for full-length and truncated αS seeds are shown. The minor increase in diffusion time observed reflects competitive binding between the small percentage of fluorescently labeled tau with the excess unlabeled. Mean $\tau_{D,\ norm}$ and SD calculated for a minimum of three measurements with αS seeds. %diff =[$\tau_D$(+αS) - $\tau_D$(-αS)]/$\tau_D$(-αS).
